# Heterozygote advantage cannot explain MHC diversity, but MHC diversity can explain heterozygote advantage

**DOI:** 10.1101/2025.05.27.656382

**Authors:** Joshua L. Cherry

**Affiliations:** Computational Biology Branch, Division of Intramural Research, National Library of Medicine, National Institutes of Health, 8600 Rockville Pike, Bethesda, MD 20894, USA

## Abstract

Several theoretical studies have concluded that heterozygote advantage makes at most a minor contribution to MHC diversity. Siljestam and Rueffler (2024) recently presented models in which heterozygote advantage alone can lead to realistically high diversity. Here I argue that heterozygote advantage cannot by itself explain MHC diversity, and that its contribution to diversity is unlikely to be large in most species. I first show that the high diversity reported by Siljestam and Rueffler is so sensitive to parameter values that the underlying phenomenon cannot explain the widespread diversity of MHC genes. I then consider a fundamental problem with explaining MHC diversity by heterozygote advantage alone: selective forces that favored heterozygotes would lead to the evolution of haplotypes having much higher fitness when homozygous, diminishing or eliminating heterozygote advantage. Diversity maintained by another force, however, might bring about adaptation to the more common heterozygous state at the expense of homozygous fitness. Thus, substantial heterozygote advantage may arise as a consequence of MHC diversity.

## Introduction

The MHC class I and II genes are the most polymorphic loci in jawed vertebrates. These genes encode proteins that present peptides, derived from host or pathogen proteins, on the surface of host cells, where they may be recognized by T cell receptors having appropriate specificities. This recognition plays several important roles in the immune system’s discrimination between self and nonself. Because MHC alleles differ with respect to the profile of peptides presented, MHC genotype can affect many aspects of immune function, including pathogen resistance and susceptibility to autoimmunity.

Various types of selection, which are not mutually exclusive, have been proposed to explain MHC diversity (reviewed in Apanius et al. (1997) and Radwan et al. (2020)). The most often discussed hypotheses involve selection for resistance to pathogens. Genotypes bearing rare alleles may be at an advantage because they share fewer pathogen sensitivities with conspecific sources of infection. Heterozygote advantage due to superior pathogen resistance has also been proposed to explain MHC diversity (Doherty and Zinkernagel, 1975).

MHC heterozygotes are thought to have greater resistance to pathogens because they present a wider variety of peptides. However, presentation of too great a variety of peptides is likely deleterious; otherwise, MHC restriction would not have evolved. The main candidates for harmful effects involve self/nonself discrimination: the elimination of more of the T cell repertoire due to reaction with self peptides and increased autoimmunity (reviewed in Woelfing et al. (2008)).

Direct empirical tests of MHC heterozygote advantage have yielded conflicting results. Some studies have found heterozygote advantage (Penn et al., 2002; Savage and Zamudio, 2011), but homozygote advantage has also been observed (Ilmonen et al., 2007).

Simulation results and theoretical considerations have suggested that heterozygote advantage makes at most a minor contribution to MHC diversity. Lewontin et al. (1978) concluded that maintenance of large numbers of alleles by heterozygote advantage is unlikely at any locus because it requires that different types of heterozygotes have very similar fitness. De Boer et al. (2004) considered MHC loci specifically and drew a similar conclusion.

Siljestam and Rueffler (2024) recently presented models that can generate realistically high MHC diversity as a result of heterozygote advantage alone. These models incorporate features that allow most heterozygous pairings of a large number of alleles to have similar, high fitness while all homozygotes have much lower fitness. These features include a saturating function relating “condition” to fitness and the multiplication across pathogens of allele averages of the efficiencies of protection. On the basis of these results, Siljestam and Rueffler argue for reconsideration of heterozygote advantage as an important force contributing to MHC diversity, though they do not claim that it is the only such force.

Here I make the case that heterozygote advantage is not by itself a tenable explanation for MHC diversity, and that its contribution to diversity is unlikely to be large. I show that, contrary to appearances, the phenomenon that leads to high diversity in the simulations of Siljestam and Rueffler depends on finely tuned parameter values, and hence is unlikely to operate in a wide variety of species. In addition, I contend that strong heterozygote advantage is not expected in the absence of diversity maintained by some other force.

The argument for the second proposition can be summarized as follows. If homozygotes were much less fit than heterozygotes because they present too limited a variety of peptides, and no other force acted to maintain diversity, MHC haplotypes would simply evolve to present a wider variety of peptides, diminishing or eliminating heterozygote advantage. Diversity maintained by another force, however, might disfavor haplotypes that can form fit homozygotes because they are less fit in the more common heterozygous state, leading to heterozygote advantage.

Consider the situation illustrated in Figure 1A, in which two (or more) MHC alleles are found at non-negligible frequencies and homozygotes suffer a fitness deficit. A homozygote for a haplotype bearing both alleles (Figure 1B), perhaps with expression of each gene reduced by a factor of two, would be about as fit as the heterozygote. A population monomorphic for this haplotype would have higher average fitness than the polymorphic population in Figure 1A because it is free of unfit homozygotes. For a polymorphic population with many alleles, this would be true for at least some allele pairs, including the pair forming the most fit heterozygote. If selection strongly favored even greater presentation breadth than provided by two alleles, expansion could continue until such selection became weak or nonexistent.

**Figure 1.**
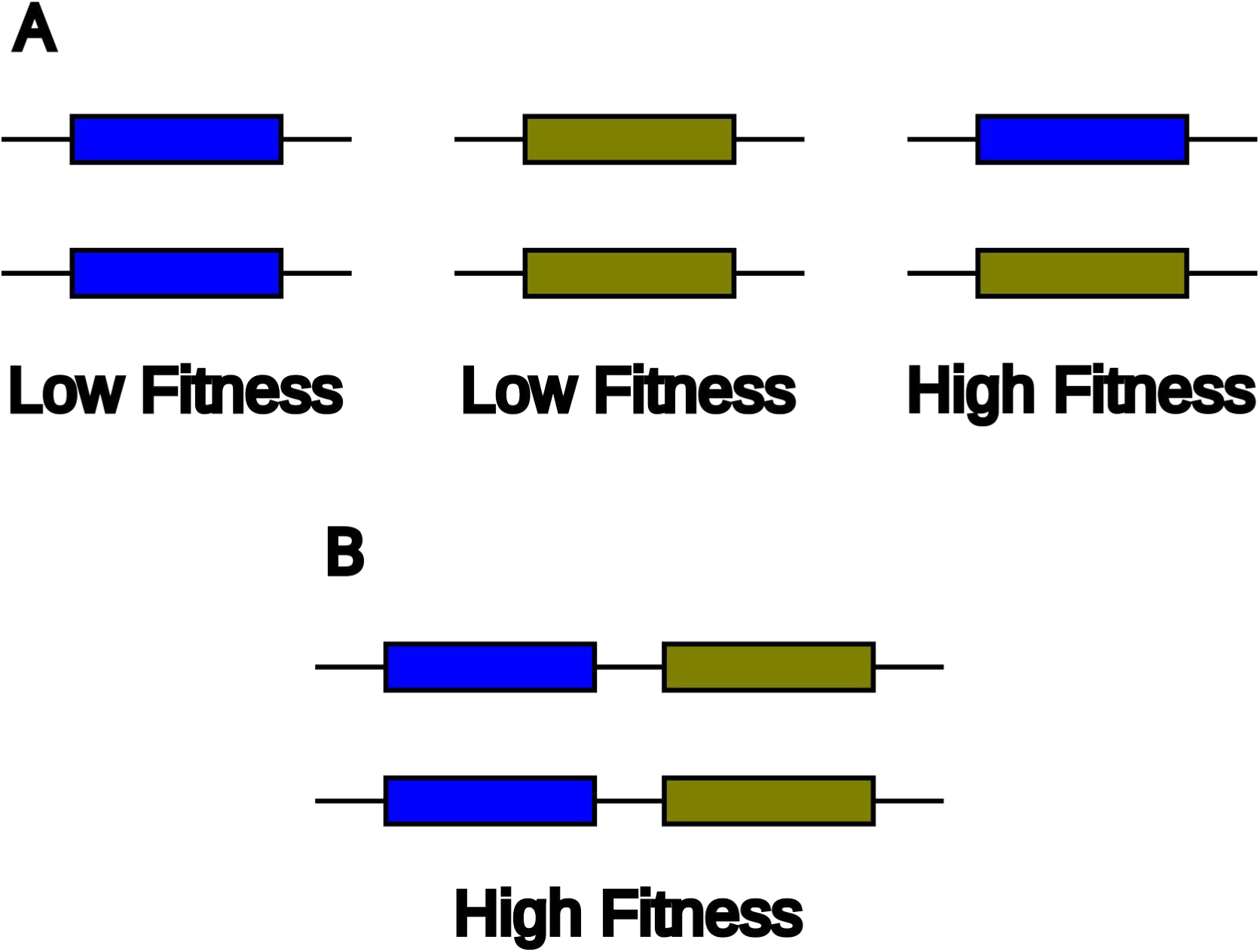
(A) A population with polymorphism and heterozygote advantage. The population includes high-fitness heterozygotes and low-fitness homozygotes. (B) A haplotype bearing two alleles in tandem forms high-fitness homozygotes. A population monomorphic for such a haplotype consists of only high-fitness individuals.

MHC gene family expansion has in fact occurred: most species carry two or more class I alpha genes and two or more class II alpha/beta pairs. This does not preclude further expansion in response to selection. In fact it demonstrates the feasibility of expansion on the necessary timescale and may facilitate further expansion through unequal recombination (Schimenti, 1999). MHC gene family expansion in response to selection has also been observed in simulations (Bentkowski and Radwan, 2019).

Another possible route to fit homozygotes is for individual MHC proteins to evolve to present a wider variety of peptides. Indeed, it is unclear why, prior to diversification, they would have evolved to be so selective that homozygotes were unfit.

After showing that the results of Siljestam and Rueffler are more sensitive to parameter values than they appear to be, I use simulations to study extensions of these models that allow the presentation breadth conferred by a haplotype to evolve. These extensions can lead to breadth expansion that diminishes heterozygote advantage and eliminates diversity.

## Results

### Sensitivity of the models of Siljestam and Rueffler to parameter values

The simulations of Siljestam and Rueffler appear to indicate that high diversity does not depend strongly on parameter values. However, these simulations varied one of the parameters, *c*_max_, in a way that compensates for changes to other parameters. As *c*_max_ is not expected to change in this way in nature (see Discussion), I assess the sensitivity of the results to parameter values in the absence of compensatory changes to *c*_max_.

I first consider the symmetric Gaussian model analyzed by Siljestam and Rueffler, with the following parameters: *N* (population size) equal to 10^5^, *m* (number of pathogens) equal to 8, *v* (“virulence”) equal to 20, *K* (half-saturation constant) equal to 1, and μ (mutation rate) equal to 5·10^-7^. With *c*_max_ determined as in Siljestam and Rueffler, these parameters lead to high MHC diversity (Figure 2, red curve), as Siljestam and Rueffler demonstrated.

**Figure 2.**
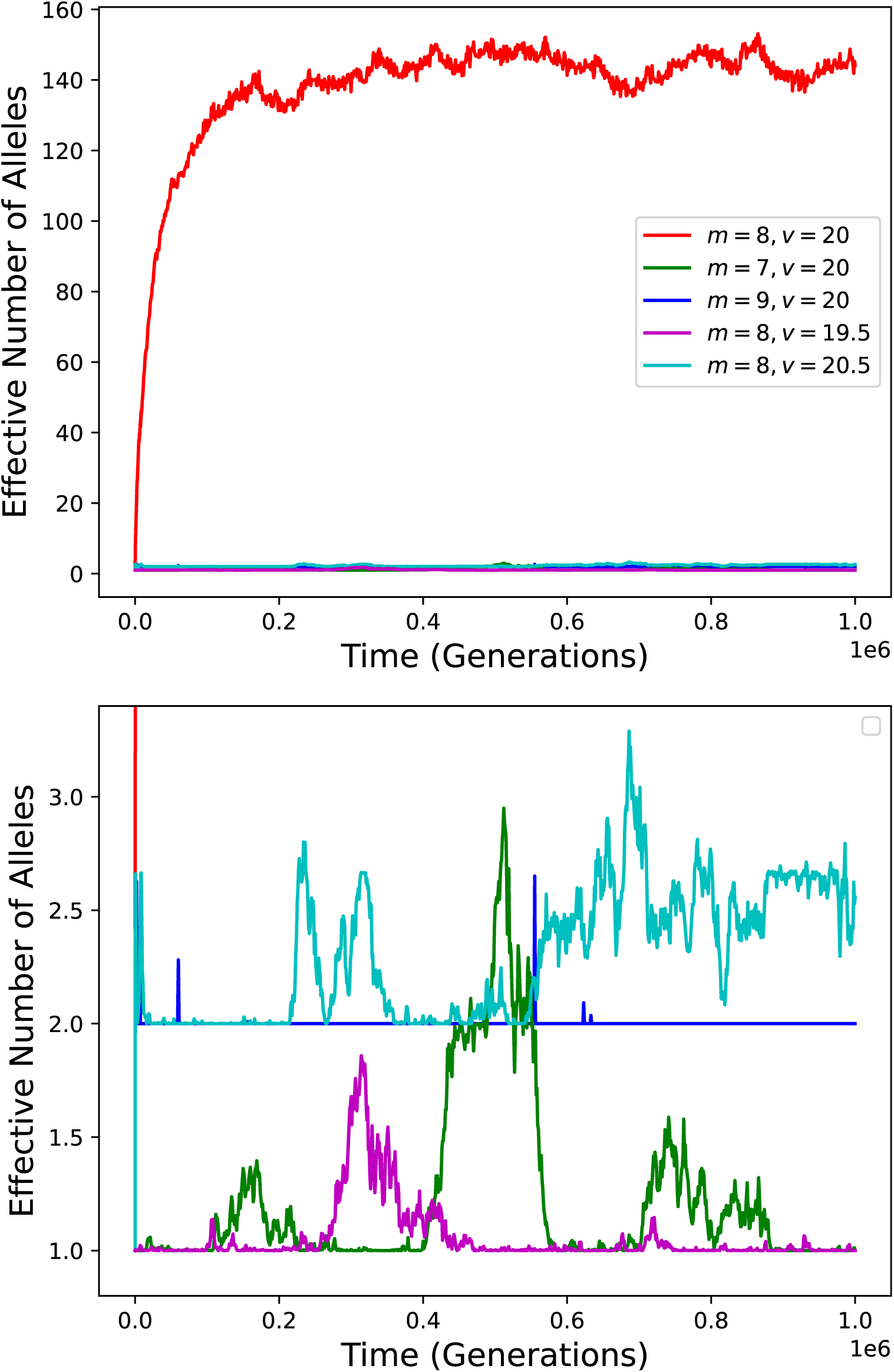
MHC diversity over time in simulations with parameter values that lead to high diversity (red curve) and with slightly altered parameter values but no compensatory adjustment of *c*_max_. The bottom panel is a vertical zoom of the top panel.

If the number of pathogens, *m*, is decreased from 8 to 7, and the other parameters, including *c*_max_, remain unchanged, diversity is mostly eliminated (Fig. 2, green curve). This is due to the fact that removal of just one pathogen increases the maximum condition achievable by a homozygote enormously, from 1 to 2.7·10^43^. The corresponding survival probability increases from 0.5 to very nearly 1 (approximately 1-3.7·10^-44^), so that no heterozygote can be substantially more fit than all homozygotes.

Note that the procedure employed by Siljestam and Rueffler would decrease the value of *c*_max_ by a factor of more that 10^43^ in response to the decrease in *m*. This change, which would not be expected upon elimination of a pathogen, would qualitatively negate the effect of changing *m*, reducing conditions of homozygotes to 1 or less.

Diversity is also largely eliminated if *m* is increased from 8 to 9 (Fig. 2, blue curve). Because condition values are decreased by the additional pathogen, fitness becomes more sensitive to condition. This results in substantial fitness differences between heterozygotes, which are not conducive to the maintenance of high diversity (Lewontin et al., 1978). During much of the simulation the population is dominated by a pair of alleles that form heterozygotes with especially high fitness.

Small changes to the parameter *v* have similar effects. Changing *v* from 20 to either 19.5 or 20.5 mostly eliminates long-term diversity (Figure2, magenta and cyan curves).

These results were produced by my own simulation code. To verify my interpretation of the model and implementation of the simulation, I have confirmed the results using a slightly modified version of the simulation code of Siljestam and Rueffler. Supplementary Text S1 describes how this verification can be done using their publicly available code. In addition, Supplementary Text S2 provides a theoretical derivation of the factor of 2.7·10^43^ mentioned above.

Siljestam and Rueffler also considered a bitstring model, which represents MHC alleles and pathogen peptides with sequences of bits rather than abstract vectors. This model is similarly sensitive to parameter values, as illustrated in Supplementary Figure S1. With *N=*10^5^, *m*=100, and *v*=9, *K*=1, μ=5·10^-6^, five peptides per pathogen, and *c*_max_ set as in Siljestam and Rueffler, equilibrium MHC diversity is high (Supplementary Figure S1A). Because equilibrium diversity varies with the randomly chosen pathogen peptides, results of many simulation runs are shown. If ten pathogens are added (i.e., *m* is raised to 110) and the other parameters (including *c*_max_) remain the same, diversity becomes much lower (Supplementary Figure S1B): the harmonic mean diversity among simulation runs (black curve) is not much above the neutral expectation of 3. If five of the 100 pathogens are removed, even lower diversity results (Supplementary Figure S1C). Equilibrium diversity also becomes very low if *v* is changed from 9 to 9.5 or 8.5 (Supplementary Figure S1D and E).

### Gene family expansion eliminates diversity in the Gaussian model

To allow the formation of haplotypes bearing multiple alleles in tandem (Figure 1B), I introduce a low rate (10^-9^ per gamete) of events in which the MHC alleles carried by both parents are combined into a single haplotype, as if by unequal recombination. I also include gene deletion at a rate of 10^-7^ per copy for multi-gene haplotypes. To evaluate the fitness of genotypes bearing more than two alleles in total, I average the pathogen-specific efficiencies of all alleles carried by a genotypes, just as Siljestam and Rueffler did for the two alleles carried by a heterozygote. The population is initially monomorphic for the optimal single-gene haplotype, as in Figure 2. Because the effective number of alleles is ill-defined for a population in which haplotypes carry different numbers of gene copies, I track the effective number of haplotypes, defined analogously. This will never be smaller than the effective number of alleles at any locus when the latter is meaningful, as haplotypes that differ at any locus are considered to be different.

The top panel of Figure 3 shows results for twenty simulation runs. Initially, single-gene haplotypes predominate and diversity begins to rise as, in Figure 2 (red curve), due to heterozygote advantage. After some time, diversity plummets, coincident with the rise of a two-gene haplotype to high frequency. Subsequently, diversity remains very low. The time that it takes for a two-gene haplotype to become established varies stochastically. In many cases it happens very quickly, before substantial diversity can develop.

**Figure 3.**
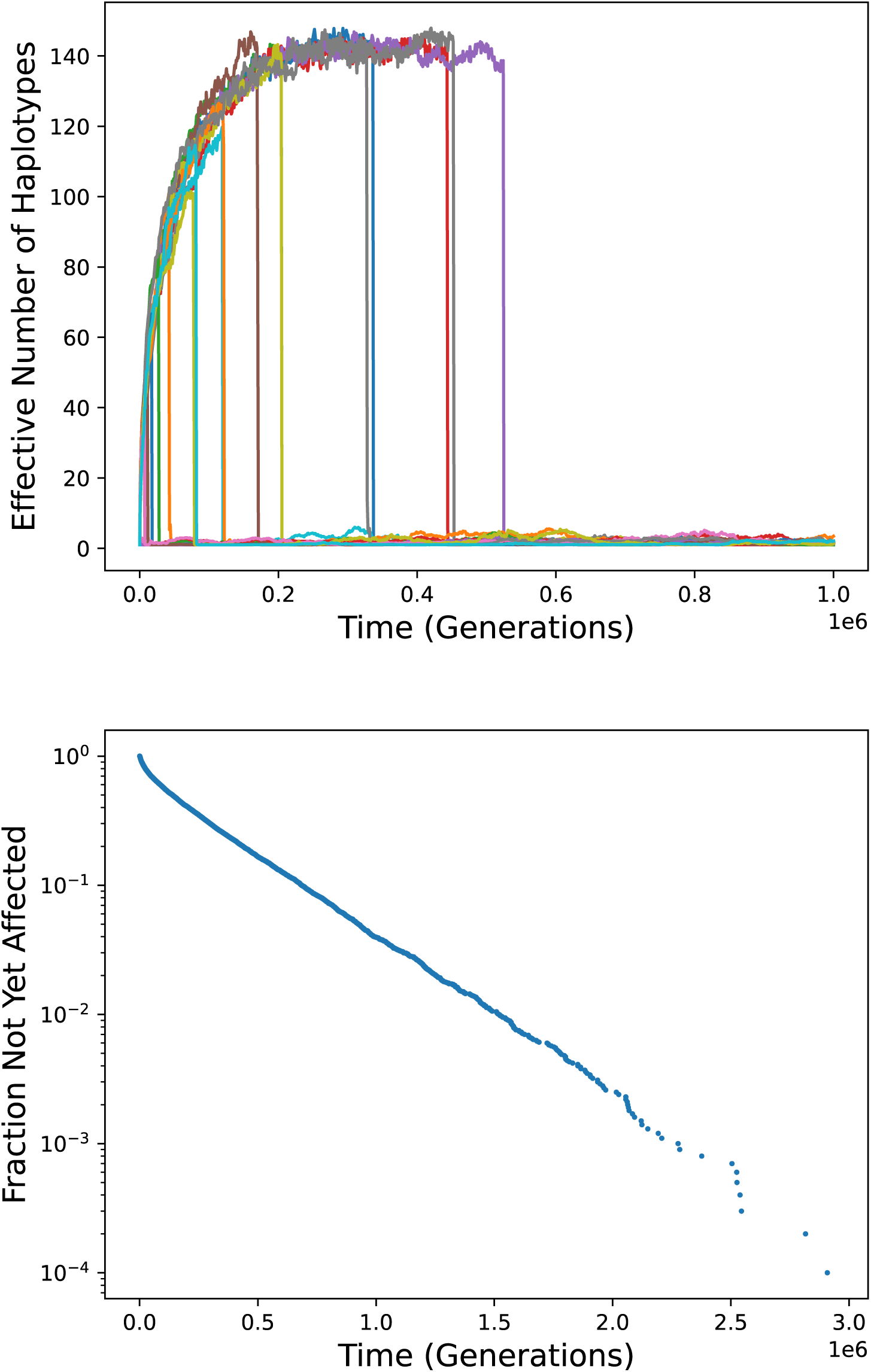
Gene family expansion in the Gaussian model. Top: diversity over time in twenty simulation runs. In each run, haplotypes bearing two alleles come to predominate, leading to a sudden loss of any diversity that has accumulated. Bottom: the distribution of time until such an event in 10,000 simulation runs.

The bottom panel of Figure 3 depicts the timing of establishment of an expanded haplotype and consequent elimination of diversity in 10,000 simulation runs. The median waiting time for such an event is about 139,000 generations. 96% of the waiting times are less than a million generations, and all 10,000 are less than three million generations. The rate of events is higher at early times because diversity is low, so that homozyosity is higher, giving and expanded haplotype has a greater advantage. Over time the rate decreases, approaching a constant value as diversity approaches an equilibrium, so that the points fall approximately along a straight line in the semi-logarithmic plot. The rate at long times is about 2.68·10^-6^/generation, or one per 3.74·10^-5^ generations. A theoretical explanation for this rate is given in Supplementary Text S1.

The survival probability of homozygotes for an established two-gene haplotype is typically greater than 0.99999, leaving little room for improvement: selection coefficients (*s*) for more fit genotypes must be less than 10^-5^, making the product *Ns* less than 1. This explains the lack of long-term diversity: a heterozygote may be more fit, but not by enough to maintain diversity in the face of genetic drift. It also explains why haplotypes bearing more than two alleles did not become established in any of the runs, despite the fact that further expansion could increase fitness.

If the parameter *K* is made very large, further expansion does occur. In some simulation runs the result is a haplotype carrying eight genes, each adapted to one of the eight pathogens. Large values of *K*, however, are not of much interest because they lead to low diversity even if haplotype expansion is not allowed.

### Gene family expansion eliminates diversity in the bitstring model

Figure 4 shows the results of adding gene family expansion to the bitstring model with parameters that lead to high diversity in its absence (Supplementary Figure S1A). The rate of haplotype expansion was again 10^-9^.

**Figure 4.**
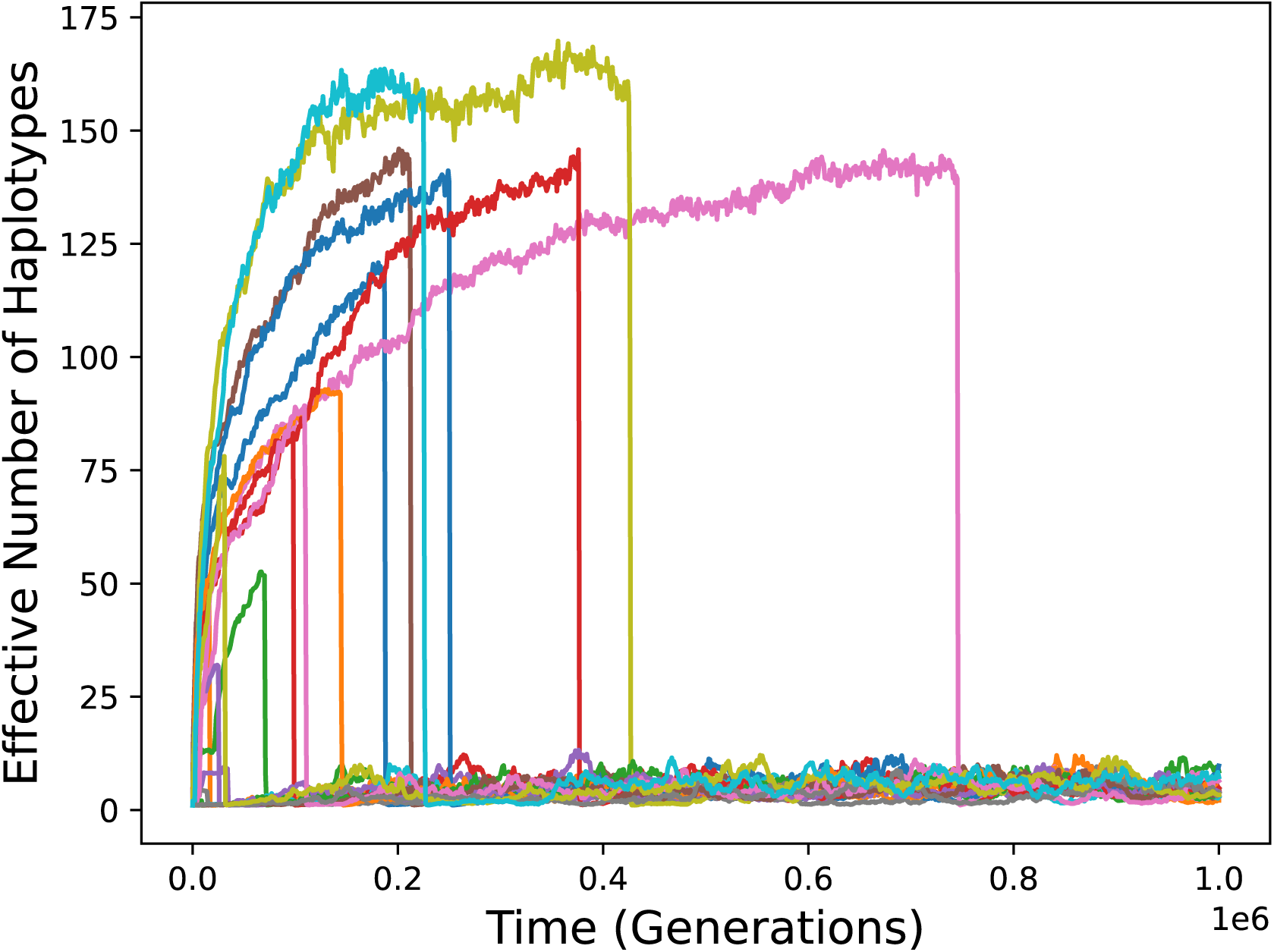
Diversity over time in simulations of the bitstring model with gene family expansion. Twenty simulation runs are shown. Diversity becomes and remains low when expanded haplotypes come to predominate.

Results are qualitatively the same as for the Gaussian model. After at most a brief period of high diversity, diversity becomes and remains low subsequent to establishment of a haplotype carrying two alleles. There is more variation among simulation runs than with the Gaussian model because runs differ with regard to the randomly chosen pathogen peptides and initial allele. Long-term diversity is larger than in the Gaussian model simulations because the mutation rate was taken to be tenfold higher, as in Siljestam and Rueffler.

### A weakly expressed additional allele is sufficient to eliminate diversity

I next consider a model in which any gene copies beyond the first carried by a haplotype are only weakly expressed. Although this model may be interpreted literally, it serves as a device for modeling small changes to presentation breadth that are achieved by any mechanism. To evaluate the fitness of genotypes that include weakly expressed alleles, I extend the averaging rule to a weighted average, with weights proportional to expression levels.

Figure 5 shows results for simulations of the Gaussian (top) and bitstring (bottom) models in which the relative expression level of additional alleles is only 2%. Parameter values are otherwise as above. The results are similar to those in which full expression was assumed (Figure 3 and Figure 4). Diversity, if it develops at all, is transient, though it tends to last somewhat longer. Haplotypes bearing two alleles (and no more), capable of forming fit homozygotes, come to predominate.

**Figure 5.**
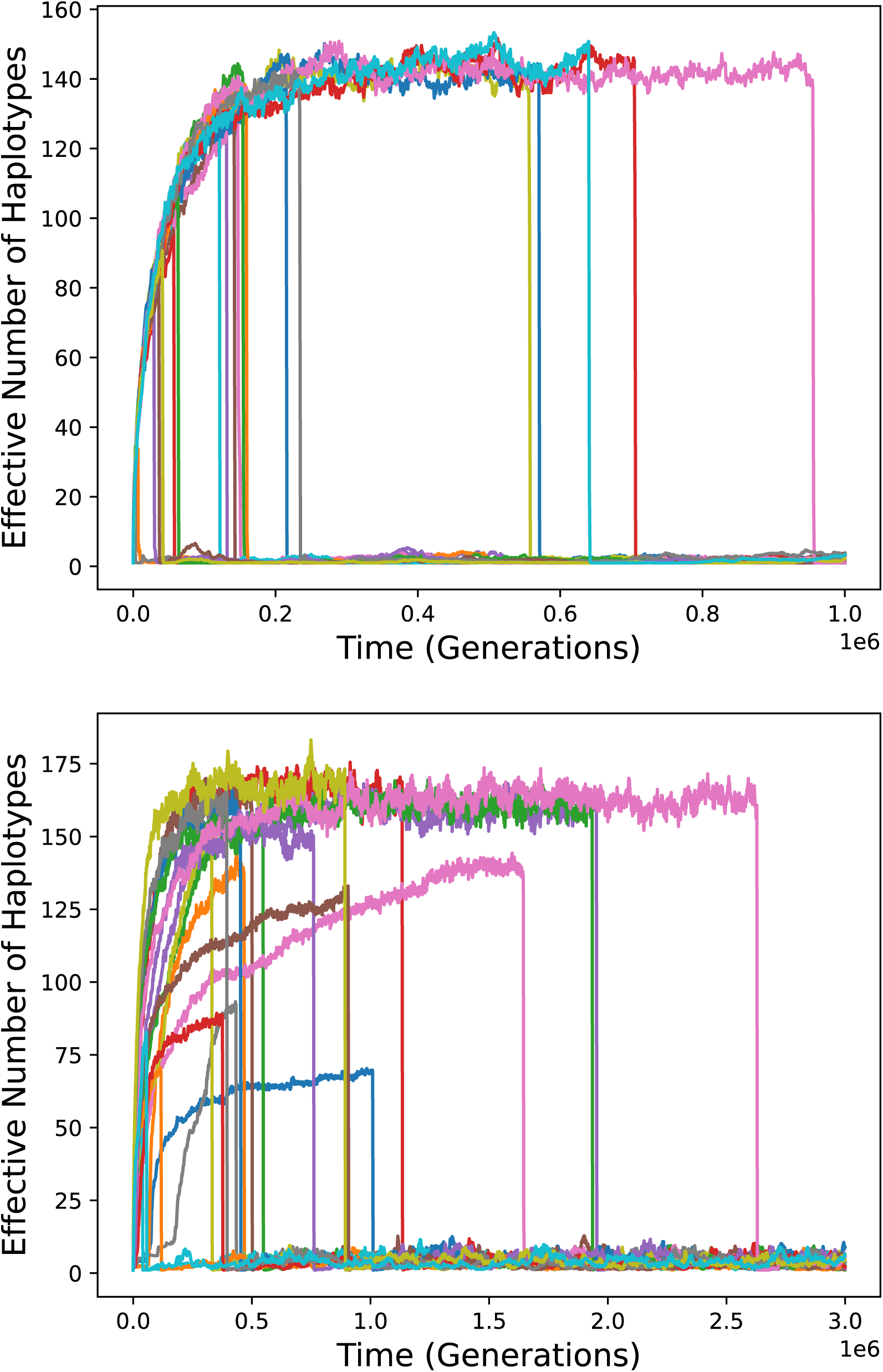
Diversity over time in simulations with gene family expansion but only weak expression of additional alleles carried by a haplotype. Top: Gaussian model. Bottom: bitstring model.

### Evolution of less restrictive MHC proteins eliminates diversity

Another possible route to greater presentation breadth is the evolution of MHC proteins that individually present a wider variety of peptides. In the bitstring model of Siljestam and Rueffler, the breadth of peptide detection conferred by any MHC allele is controlled by the model parameter *v*. Siljestam and Rueffler interpret *v* as pathogen virulence, presumably because more selective detection can, in their model, only decrease the effectiveness of the immune system. Since, however, the breadth of detection is determined in part by the MHC protein sequence, I allow *v* to be a mutable property of MHC alleles, controlled by an additional bit that mutates at the same frequency as the other bits in the model. Thus, one in seventeen MHC mutations changes the breadth of presentation rather than the optimal matching peptide.

For the initial allele, *v*=9 as in Figure 4. Mutation can change this to a lower value, which corresponds to a wider presentation breadth. Subsequent mutation can change it back to 9. The same value of *c*_max_, computed assuming *v*=9, applies to all alleles, as it would be inappropriate to apply different *c*_max_ values to different alleles in the same population given the biological meaning of this parameter (see Discussion).

It is conservative in this context to assume that increased presentation of some peptides comes at the expense of reduced presentation of others. Therefore, when increasing presentation breadth by lowering *v*, I also reduce the detection probability for every peptide by a constant factor, such that the mean detection probability across all 2^16^ possible peptides remains constant.

Figure 6, top, shows simulation results for this model when the alternative value of *v* that an allele can specify is 8.12. A change from *v*=9 to *v*=8.12 increases the effective number of peptides presented (see Methods) by nearly a factor of two, and thus approximates the increase in presentation breadth that comes from increasing the number of alleles from one to two when the alleles are dissimilar. In one hundred simulation runs, the effective number of alleles never rises above 18.5, and its harmonic mean among runs is below the neutral expectation for the length of the simulations. Alleles with increased presentation breadth come to predominate very quickly: this occurs by generation 5300 in every simulation run. The evolution of fit homozygotes and the concomitant elimination of any diversity is much faster than in Figure 4 because the rate of breadth-increasing mutation is assumed to be much higher than the rate of gene family expansion.

**Figure 6.**
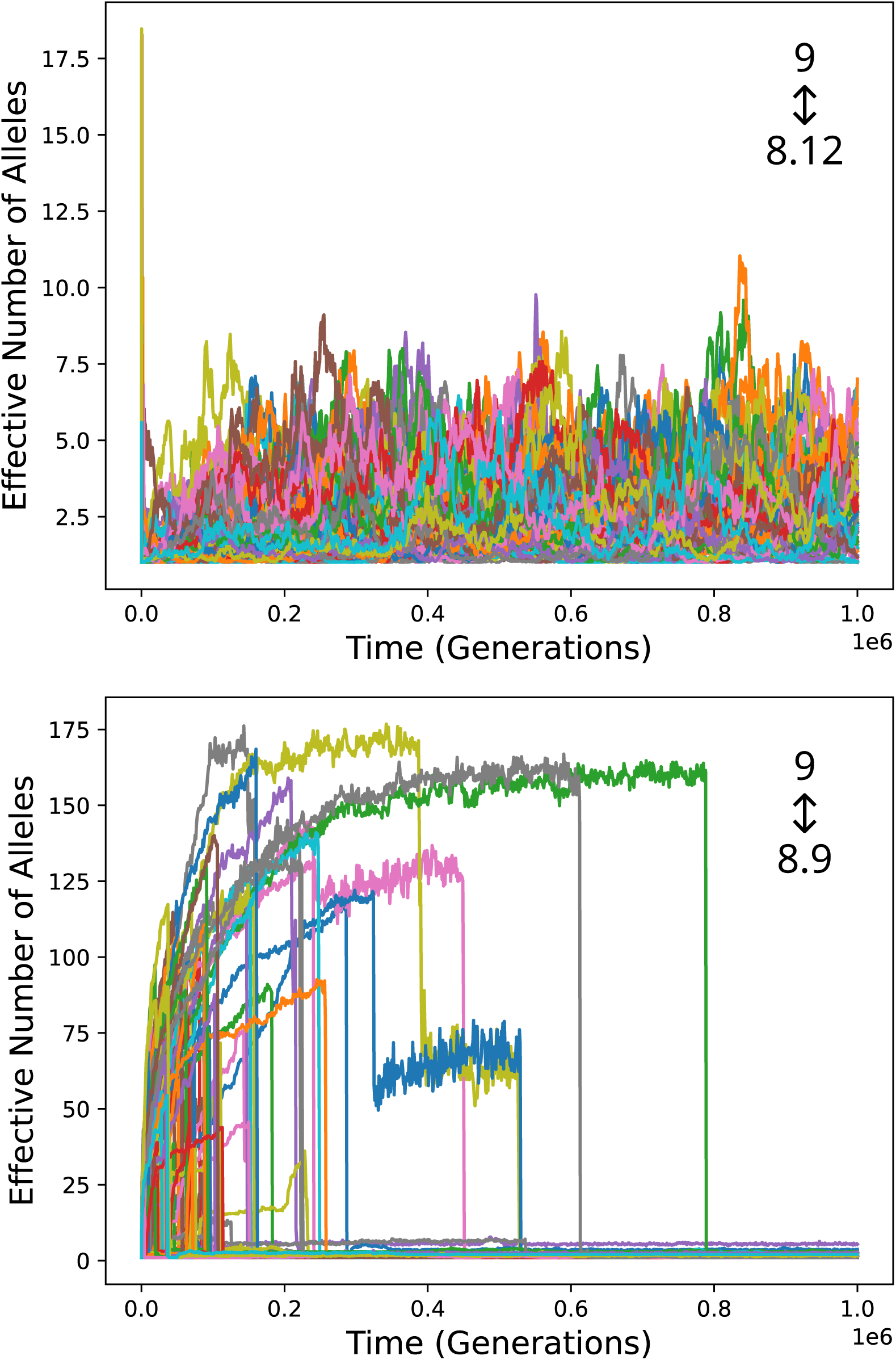
Simulation results for a model in which mutation can alter the breadth of peptides presented. The initial allele specifies *v*=9, and *c*_max_ is set accordingly. Mutation can change *v* to 8.12 (top) or 8.9 (bottom). Results of one hundred simulation runs are shown in each plot.

Additional simulations confirm that diversity is quickly eliminated even if it is allowed to rise to equilibrium values before breadth-increasing mutations are allowed (Supplementary Figure S2). These results illustrate another means by which the variety of peptides presented can be increased without allelic diversity and the burden of low-fitness homozygotes.

Figure 6, bottom, shows that diversity is still eliminated, although more slowly, when mutation can only bring about a more modest increase in presentation breadth: a change from *v*=9 to *v*=8.9, which corresponds to an 8% increase in breadth. In many cases there is a substantial initial rise in diversity because the initial allele confers low fitness in the homozygous state. Over time, an allele capable of forming fit homozygotes evolves and comes to predominate, mostly eliminating heterozygote advantage and allelic diversity.

### Incorporating a cost of overly broad presentation

The simulations presented so far do not include any explicit deleterious effects of overly broad presentation of peptides. This is not an issue for the homozygous genotypes that come to predominate, which have presentation breadths similar to or narrower than those of heterozygotes with unexpanded presentation breadths. However, when rare, a haplotype with increased presentation breadth occurs mainly in heterozygotes having greater total presentation breadth than “ordinary” heterozygotes. Low fitness of these intermediates might constitute a barrier to the establishment of a haplotype with larger presentation breadth and the consequent elimination of diversity.

I first consider a model of what may seem like a severe cost of overly broad presentation: the detection probability of any peptide due to any allele is cut in half. Supplementary Figure S3 shows results for simulations of the bitstring model with *v* evolvable from 9 to 8.12 and such a cost.

Despite this cost, diversity is eliminated fairly quickly. Due to the saturating fitness function, this cost has little effect on fitness for many genotypes, and the establishment of a haplotype forming fit homozygotes is not much delayed.

A substantial cost of overly broad presentation can be imposed directly. With a 5% fitness cost, a persistent low diversity state is quickly established in 9951 of 10,000 simulation runs (99.5%). If zero fitness is assigned to all diploid genotypes with expanded presentation breadth, the largest possible harmful effect of overly broad presentation, the outcome depends strongly on the initial state (Figure 7). If the initial allele has *v* equal to 9, high diversity develops and persists in 81 of 100 simulation runs (Figure 7, top). However, if the initial allele has *v* equal to 8.12, diversity remains very low (Figure 7, bottom). It should be noted that a breadth of presentation corresponding to *v*=9 would be severely maladaptive in a monomorphic population, making the first initial condition difficult to explain.

**Figure 7.**
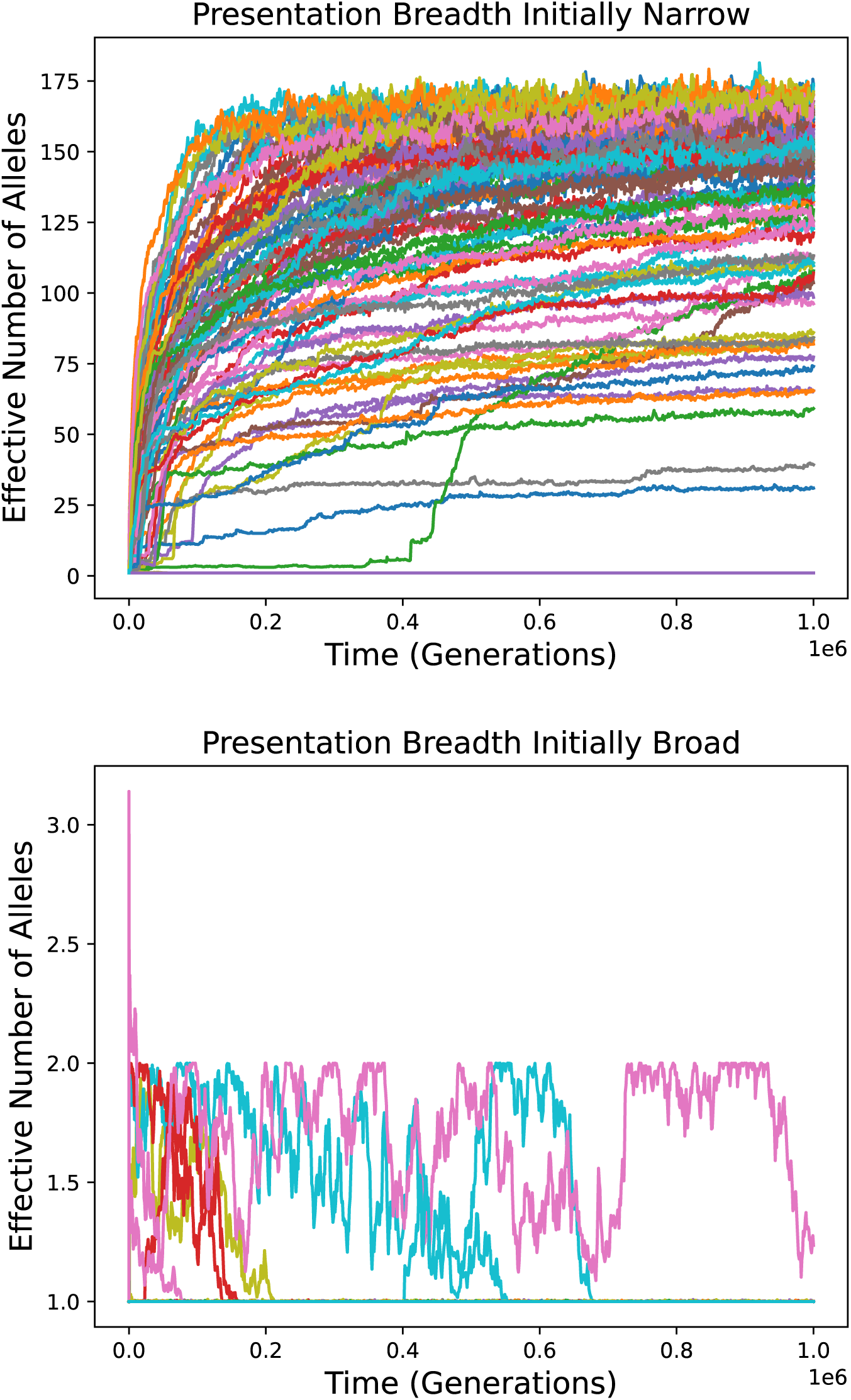
Diversity over time in simulations of the bitstring model in which *v* can evolve between 9 and 8.12 and overly wide presentation breadth is lethal. The initial allele has narrow presentation breadth (*v*=9) in the top panel and broader presentation breadth (*v*=8.12) in the bottom panel. One hundred simulation runs are shown for each. Note that many curves in both plots fall on top of each other, with effective number of alleles close to 1 throughout the simulations.

## Discussion

### High diversity requires fine tuning of parameters

A model predicting high MHC diversity for only a narrow region of parameter space cannot explain universally high diversity, as parameters can be expected to vary over evolutionary time. As illustrated in Figure 1 and Supplementary Figure S1, the high MHC diversity produced by the models of Siljestam and Rueffler is highly dependent on parameter values. For example, increasing or decreasing the number of pathogens slightly can eliminate most of the diversity. This sensitivity is not evident in the results of Siljestam and Rueffler due to the adjustment of one parameter, *c*_max_, so as to compensate for differences in others. The value of *c*_max_ is not expected to change with factors such as the number of pathogens as it represents the “condition” of an individual unaffected by pathogens.

This difficulty is not specific to the details of the models of Siljestam and Rueffler, but is inherent in the phenomenon invoked to allow high diversity. Central to this phenomenon is a saturating (or at least downwardly concave) relationship between an individual’s condition and its fitness. A helpful simplification is to think of the fitness function as sensitive to condition for values below some threshold and insensitive to condition for values above it. A requirement for high diversity is that the conditions of homozygotes and heterozygotes lie mainly on opposite sides of this divide. If, for any reason, condition values were to become generally higher, so that homozygotes joined heterozygotes in the insensitive regime, there would be little fitness difference between homozygotes and heterozygotes and hence little diversity. If condition values were to become lower, moving heterozygote conditions into the sensitive range, fitness would differ substantially among heterozygotes and diversity would be low for the reason put forth by Lewontin et al. (1978).

The biological implication is that any change that affects condition by as much as the difference between MHC heterozygotes and homozygotes will eliminate high equilibrium diversity. In order for the phenomenon invoked by Siljestam and Rueffler to make a large contribution to MHC diversity across species, it would have to be that changes that occur over evolutionary time—changes to the diversity and properties of pathogens, increases or decreases in food supply, the transition from aquatic to terrestrial habitat, the advent of flight—all have smaller effects on condition than does MHC heterozygosity status. As this is implausible, the phenomenon cannot explain a substantial portion of MHC diversity in most species.

### Selection can bring about elimination of heterozygote advantage

The simulation results presented in Figs. 3, 4, 5, and 6 support the suggestion that MHC diversity due to heterozygote advantage alone will not persist if, as in reality, a haplotype can evolve to present a wider variety of peptides. In some simulation runs, no substantial diversity develops; in others, diversity is high only transiently. In either case, haplotypes with increased presentation breadth come to predominate. These form fit homozygotes, so there is subsequently little heterozygote advantage and hence little diversity.

These dynamics are consistent with what is known about established cases of polymorphism maintained by heterozygote advantage. In the best-know examples, which involve globin alleles that cause sickle cell anemia and thallasemia in homozygotes (Allison, 1954; Haldane, 1949), the advantage enjoyed by heterozygotes is some degree of resistance to falciparum malaria, which has been a selective force for only about 7000 years or less (Livingstone, 1958; Sharp et al., 2020). In a review of established cases of polymorphism maintained by heterozygote advantage, Hedrick (2012) concluded that most or all are of recent origin.

Typical persistence times of diversity in the simulations ranged from thousands to hundreds of thousands of generations, which is very little time compared to the hundreds of millions of years since the origin of MHC. These times, however, depend on parameter values and the details of the model.

The rate of formation of haplotypes bearing all parental alleles in tandem, as if by unequal recombination, was taken to be 10^-9^ in the simulations that included this possibility. The rate of segmental duplication, which frequently occurs by unequal recombination, has been estimated to be about ∼10^-9^ per year per gene in vertebrates (Cotton and Page, 2005). Thus, the rate assumed in the simulation is realistic if generations are equated with years. The rate of unequal recombination in the MHC region may in reality be much larger than this due to the presence of multiple gene copies, many pseudogenes, and a high density of repetitive elements (Gaudieri et al., 1997; Kulski et al., 1999; Westerdahl et al., 2022).

In the simulations that allowed mutation to change the breadth of peptides presented by an MHC molecule, such mutations occurred at one sixteenth the rate of mutations altering the optimally bound peptide. It is reasonable to assume that the rates of the two types of change are comparable. It would be surprising, in fact, if mutations that affect peptide binding preferences do not frequently change the breadth of binding to some extent. In any case, establishment of a low-diversity state is extremely rapid in Figure 6, top, and remains reasonably fast if the rate of breadth-expanding mutation is decreased by a factor of one hundred.

Although the simulations were necessarily based on particular models, chosen because they result in especially high diversity under some conditions, most of the important assumptions are not model-specific. That a homozygote for a two-allele haplotype (Figure 1B) can mimic the phenotype of a heterozygote is expected from general considerations. The size of the advantage of forming fit homozygotes is inextricably linked to the strength of diversity-promoting heterozygote advantage.

The main point of uncertainty concerns the fitness values assigned to certain transitional genotypes in the simulations. A two-gene haplotype, when rare in a high-diversity population, occurs mainly in heterozygotes bearing a total of three alleles. These typically have high fitness according to the model, but the reality might be different because presentation of too wide a variety of peptides is expected to be detrimental. If three-allele genotypes were sufficiently unfit, this would be a barrier to the establishment of a two-allele haplotype in the presence of diversity. An analogous possibility applies to individual alleles with expanded presentation breadth.

This possibility is addressed to some extent by the results presented in Figure 5 and the bottom panel of Figure 6. These simulations show that haplotypes with only slightly increased breadth of presentation, for which any ill effects in transitional heterozygotes are expected to be small, can eliminate diversity. Furthermore, assuming that overly broad presentation interferes substantially with pathogen peptide detection has little effect on the results (Supplementary Figure S3). Although these results are specific to the models of Siljestam and Rueffler, they reflect features of the models that are necessary for high MHC diversity.

Strong selection against transitional genotypes might nonetheless impose a bidirectional barrier between population states. Adding a substantial 5% fitness cost to one model did little to prevent the establishment of alleles with increased presentation breadth when diversity is initially low, though if diversity becomes high this transition is greatly inhibited. Furthermore, if overly broad presentation is assumed to be lethal, this transition often fails to occur before sufficient diversity accumulates to inhibit it strongly (Figure 7).

A cost of overly broad presentation, it must be emphasized, would not favor the high-diversity state; it would at most slow the transition between the high-diversity state and a low-diversity state with higher fitness. Thus, invoking this cost to explain MHC diversity by heterozygote advantage alone requires accepting that most jawed vertebrates are trapped in a metastable state with a fitness deficit that would be alleviated by a simple genetic change. This state would have to have persisted, in multiple lineages, for hundreds of millions of years, during which numerous complicated adaptations emerged.

Should this scenario seem plausible, two concrete points can be made against it. First, it requires that MHC haplotypes evolved presentation breadth so narrow that homozygotes were highly unfit prior to the emergence of MHC diversity, when homozygotes would have been common. How such a maladaptive situation could have come about is unclear. Second, it is doubtful that such a state, if ever established, would in reality persist for hundreds of millions of years. Though various factors that might allow escape could be mentioned, the best reason for doubt is empirical: the copy number of MHC loci varies substantially between and within species (Figueroa et al., 2001; Chen et al., 2019; O’Connor et al., 2019; Mishra et al., 2020; Wong et al., 2022; Pyo et al., 2022; Westerdahl et al., 2022; Gaigher et al., 2023; He et al., 2023; Minias et al., 2023; He et al., 2024). Whatever the selective constraints on presentation breadth may be, they do not prevent the type of change necessary for the elimination of heterozygote advantage.

Thus, heterozygote advantage alone cannot explain MHC diversity because substantial heterozygote advantage is not expected in the absence of some other force that promotes diversity. However, strong heterozygote advantage might develop and persist as a consequence of diversity driven by some other force.

### Heterozygote advantage as a consequence of diversity

If there is an intermediate optimum for the total breadth of peptides presented by an individual, the optimum for a haplotype will be a compromise between fitness in the homozygous and heterozygous states. In the absence of some other force that drives diversity, haplotypes become well-adapted to the homozygous state, which is common due to low diversity, and there is no substantial hetrozygote advantage; the rare heterozygote might even be substantially less fit. Suppose, however, that some selective force other than heterozygote advantage drives high diversity. Then homozygotes will be rare and heterozygotes common unless inbreeding is intense. Selection will drive haplotype presentation breadth toward a lower value, closer to the optimum for the heterozygous state and further from the optimum for homozygotes. As a consequence, substantial heterozygote advantage may develop. Whether it does depends on the frequency of homozygosity and the details of the fitness landscape; the opposite effect, homozygote advantage, is also possible, particularly in highly inbred species. Any heterozygote advantage that does develop will contribute to selection for diversity. This effect, however, would be secondary to, and dependent on, diversity maintained by some other force, without which there would be no substantial heterozygote advantage.

## Methods

The model of Siljestam and Rueffler (2024), with modifications described in Results, was simulated using a program written in the Python programming language. Characteristics of the evolving population, including the effective number of alleles or haplotpes, were recorded at intervals of 1000 generations or less. The program is available at https://github.com/jlcherry/MHC_het_advant.

For assessment of sensitivity of the Gaussian model to parameter values, *c*_max_ was set to the same value for all simulations, regardless of the values of the other parameters. This value was that used by Siljestam and Rueffler for *m*=8 and *v*=20, which is exp(700) (see Supplementary Text S2). The procedure for the bitstream model was more complicated because the value used by Siljestam and Rueffler depends not only on the parameter values, but also on the peptides chosen randomly for each pathogen. When *v* was varied, *c*_max_ was calculated by the procedure of Siljestam and Rueffler, but using *v*=9 rather than the actual value of *v*. When *m* was varied, *c*_max_ was always calculated using the peptides for 100 pathogens. For *m≥*100, these were the peptides for first 100 of the *m* pathoges. For *m*<100, peptited for 100 pathogens were randomly generated and used to calculate *c*_max_, but only the first *m* pathogens were retained for the simulation. Results are quite similar if a *c*_max_ value obtained from a simulation run with *m*=100 and *v*=9 is used with other parameter values regardless of the random peptides generated.

For the purpose of assessing the time until establishment of haplotypes with increased breadth of presentation, establishment was defined as a mean number of alleles per haplotype of 1.99 or above (gene family expansion) or a frequency of alleles with the lower value of *v* greater than or equal to 0.99 (breadth expansion by mutation).

Estimation of the long-term rate of establishment of expanded haplotypes in the simulations shown in Figure 3, bottom, was based on the 396 establishment events occurring after one million generations. The waiting time for each of these events—the time between the millionth generation and the first such event, and the times between subsequent events—was multiplied by the number of simulations awaiting establishment prior to the event. The average of these products was taken as the estimate of the mean waiting time, and its reciprocal as the estimate of the rate of establishment events.

In order to quantify the effect of *v* on presentation breadth in the bitstring model, I consider the detection probabilities of all 2^16^ possible bitstrings that represent pathogen peptides. I divide these probabilities by their sum and treat the result as a probability distribution among peptides. From this, I calculate the effective number of peptides as the reciprocal of the sum of squared probabilities. This measure is analogous to the effective number of alleles in population genetics. For any *v*, this measure is the same for all alleles and need only be calculated for a single representative.

## Acknowledgements

This work was supported by the Intramural Research Program of the National Institutes of Health (NIH). The contributions of the NIH author are considered Works of the United States Government. The findings and conclusions presented in this paper are those of the author and do not necessarily reflect the views of the NIH or the U.S. Department of Health and Human Services.

## Supplementary Text S1

### Confirmation of Parameter Sensitivity with the Simulation Code of Siljestam and Rueffler

Here I describe how my conclusions about sensitivity to parameter values can be verified using the simulation code provided by Siljestam and Rueffler themselves, with only small, easily understood modifications. Using their code requires the Matlab software package.

The starting point is the Matlab file `MHC_sim_Dryad.m`, available at https://datadryad.org/dataset/doi:10.5061/dryad.69p8cz98j. First, we can add a line that prints the value of the variable logcmax, which represents the natural logarithm of cmax determined and used by the code. Below line 116 (‘prework’), add the line ‘logcmax’ (with no semicolon).

Now, at the Matlab prompt, execute `MHC_sim_Dryad(false, 8, 20, 1)` to run the simulation for the Gaussian model with *m*=8, *v*=20, and *K*=1. The output will indicate that logcmax=700, in accord with the theoretical factor exp(100*(*m*-1)) derived in Supplementary Text S2. The allelic diversity, *n*_e_, will rise to a steady state-level of about 140, as in the red curve of my Fig. 2.

Now lower *m* to 7, i.e, run `MHC_sim_Dryad(false, 7, 20, 1)`. The output will indicate that logcmax=600. This confirms that lowering *m* by 1 causes the code to lower the value of *c*_max_ by a factor exp(100)=2.7·10^43^, which must also be the factor by which the condition of the most fit homozygote would increase without this adjustment.

With the change of *m* to 7 and the compensatory change in *c*_max_, steady-state allelic diversity remains high. But what if *m* changes but *c*_max_ remains the same, as it would in reality? To find out, we can fix the value of *c*_max_ to the value used with *m*=8 by adding the following line below the line previously added: ‘logcmax = 700’. With this additional modification in place, executing `MHC_sim_Dryad(false, 7, 20, 1)` confirms that without a compensatory change to *c*_max_, lowering *m* from 8 to 7 mostly eliminates allelic diversity, in accord with the corresponding curve in my Fig. 2. Similarly, raising *m* from 8 to 9, or changing *v* from 20 to 19.5 or 20.5 (executing `MHC_sim_Dryad(false, 8, 19.5, 1)` or `MHC_sim_Dryad(false, 8, 20.5, 1)`), largely eliminates diversity, confirming the other results in my Fig. 2. Results for the bitstring model can also be confirmed, though this requires additional changes to the code.

## Supplementary Text S2

### Derivation of the Factor of 2.7·10^43^

As specified by Siljestam and Rueffler, the positions of the *m* pathogens in (*m*-1)-dimensional antigenic space correspond to the vertices of a regular simplex centered at the origin, with distance between vertices equal to 1. The squared distance from the origin to each of the *m* vertices of such a simplex is (*m*-1)/2*m* (https://polytope.miraheze.org/wiki/Simplex). Thus, the sum of the *m* squared distances is (*m*-1)/2. For the (0, 0) homozygote, condition is multiplied by a factor of exp(-(*vr*)^2^/2) for each pathogen, where *r* is the distance from the origin. It follows that, with *v*=20, all the pathogens together decrease condition by a factor of exp(20^2^·(*m*-1)/4) = exp(100·(*m*-1)). Thus, increasing or decreasing *m* by 1 changes this value by a factor of exp(100) = 2.7·10^43^.

## Supplementary Text S3

### The Rate of Establishment of Expanded Haplotypes

In simulations of the symmetric Gaussian model with gene family expansion (Results), the long-term rate at which expanded haplotypes came to predominate, eliminating diversity, was about 2.68·10^-6^/generation. This rate can be understood as follows. The average fitness of homozygotes for circulating single-gene haplotypes is only about 0.06, while most other genotypes have fitness close to 1. The selective advantage of an expanded haplotype, when rare, is therefore nearly equal to the rate of homozygosity, or about 1/*n*_e_, where *n*_e_ is the effective number of alleles. With only single-gene haplotypes, the harmonic mean of *n*_e_ is about 143 at equilibrium, so this selective advantage is about 0.00697. The fixation probability of an expanded haplotype is largely determined by its advantage when rare largely because fixation is almost guaranteed once it rises to a fairly low frequency. Thus, the fixation probability of a newly-arisen (single-copy) expanded haplotype is approximately 2*s*, or about 0.0139. Such haplotypes arise at a rate of 2*N*·10^-9^, so fixations are predicted to occur at a rate of 2.79·10^-6^/generation, very close to the observed rate of 2.68·10^-6^.

**Fig. S1.**
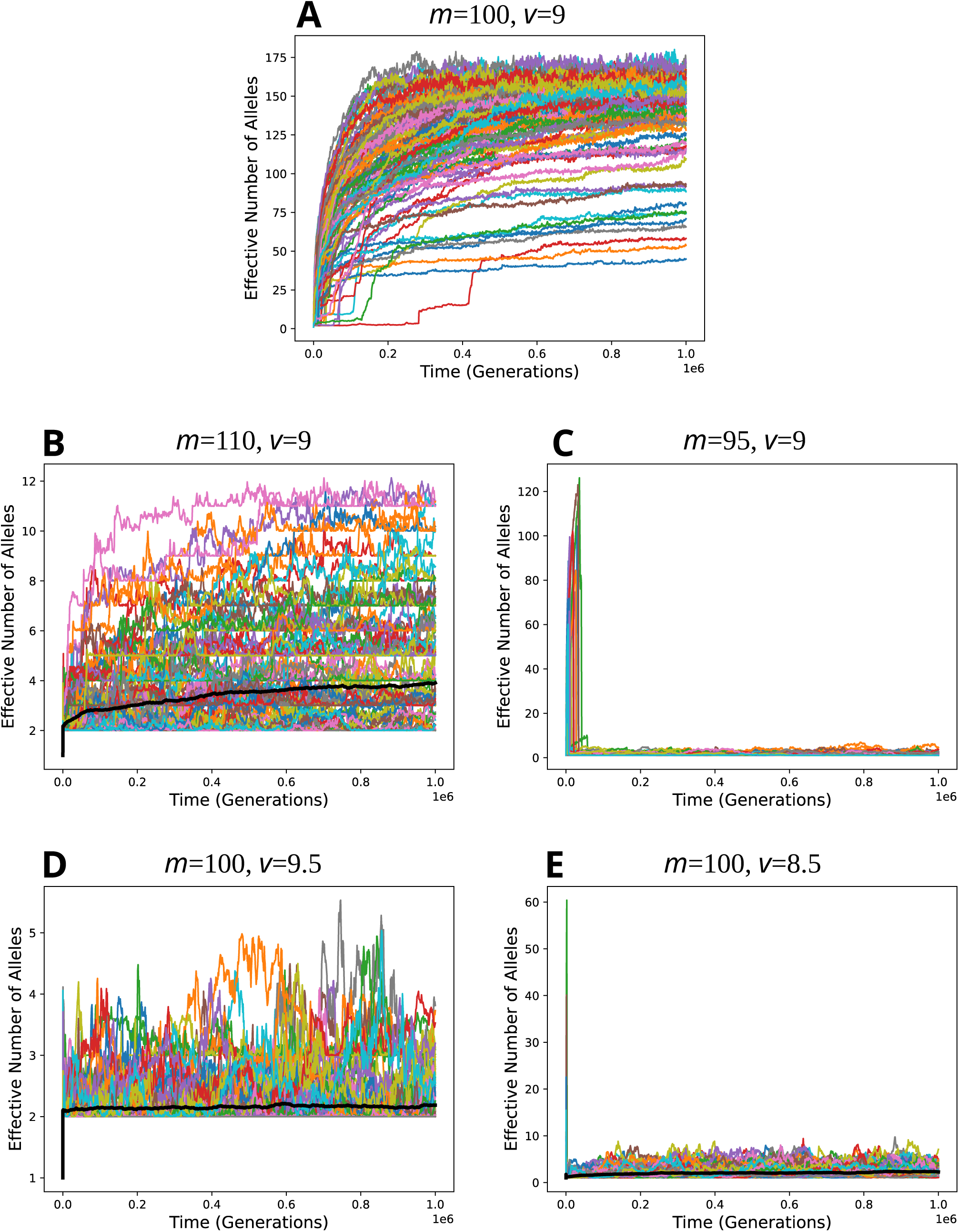
Simulation results for the bitstring model with various values of parameters *m* and *v* and identical values of other parameters, including *c*_max_. One hundred simulations runs are represented in each plot. Equilibrium diversity is high with *m*=100 and *v*=20 (**A**), but low if these parameters are changed slightly (**B-E**). The thick black curve in some of the plots (**B**, **D**, and **E**) represents the harmonic mean among runs.

**Fig. S2.**
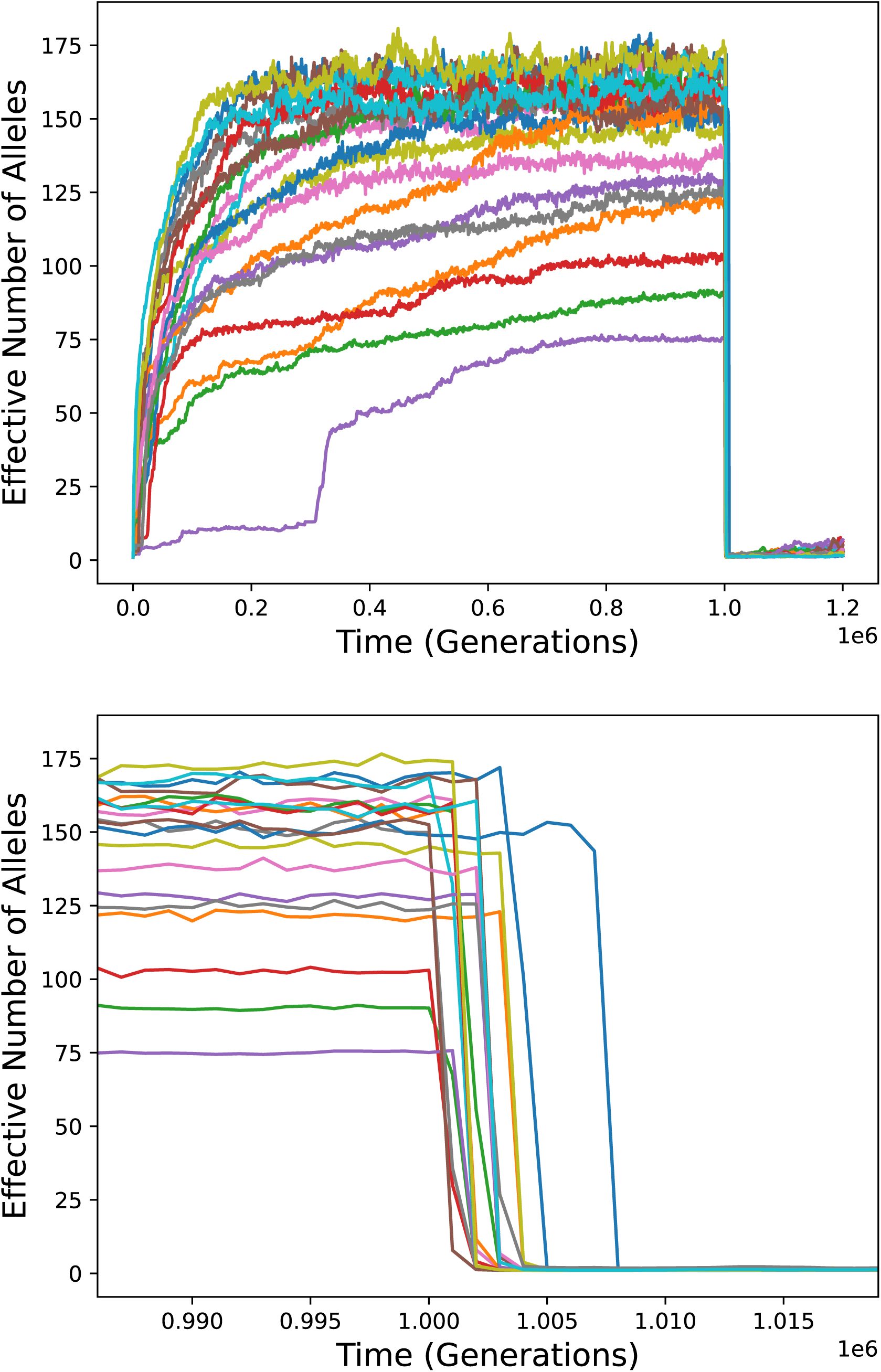
Simulations like those in Fig. 6, top, except that diversity is allowed to accumulate for one million generations before mutations affecting the breadth of presentation are allowed. In all 20 runs, diversity collapses quickly once such mutations are allowed. The bottom panel is a horizontal zoom of the top panel.

**Fig. S3.**
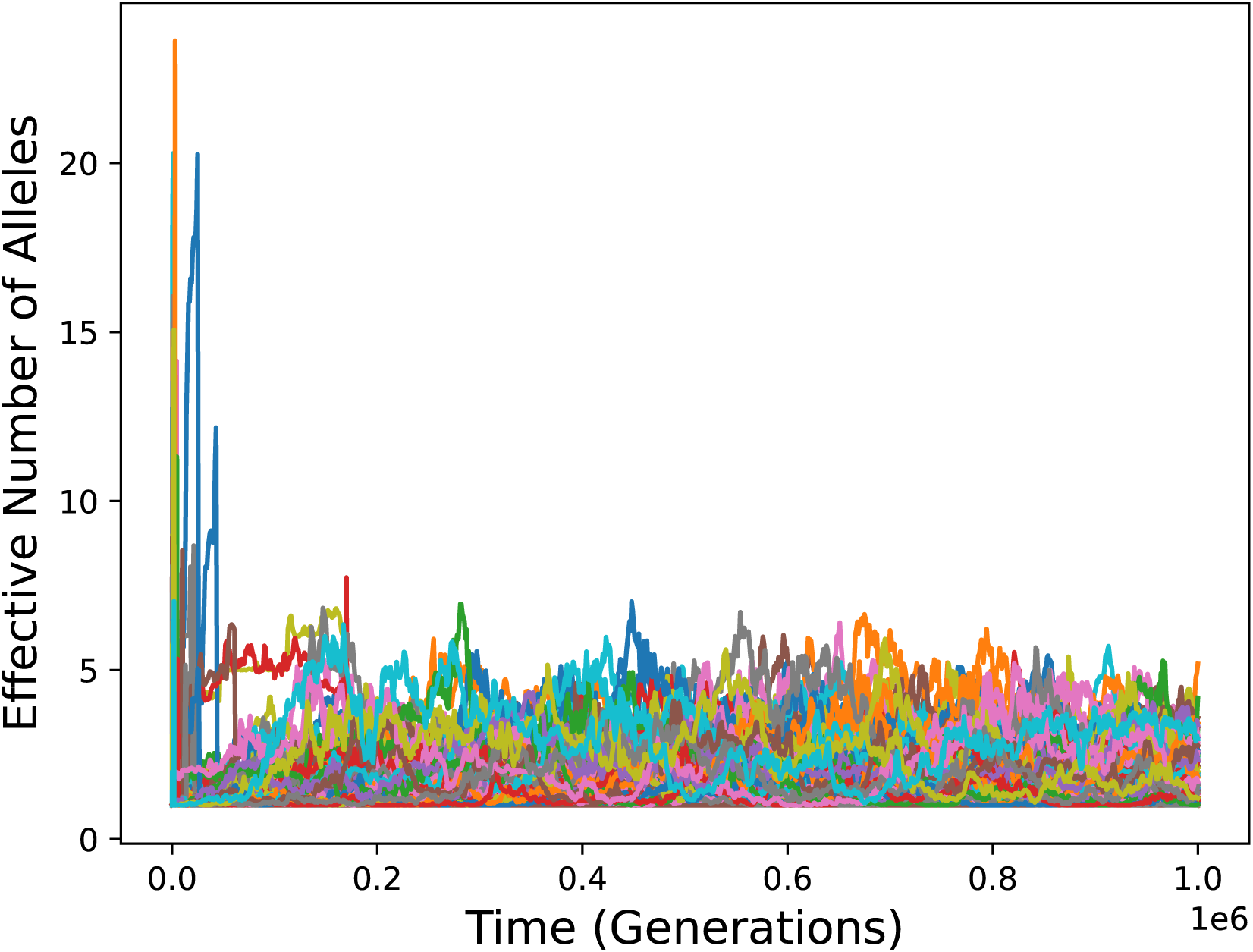
Simulations incorporating a 50% decrease in all peptide detection probabilities in individuals with certain genotypes. Other conditions are as in Fig. 6, top. Despite the inclusion of this effect, alleles with higher presentation breadth quickly come to predominate, preventing the development of high diversity.

## Notes

### Competing Interest Statement

The authors have declared no competing interest.

### Summary of Updates

Revisions based on eLife reviewer comments. These include mainly some additions to the Introduction, expansion of the end of the Discussion, addition of some details to Methods, and addition of Supplementary Text S1 and S2, which provide additional support for the conclusion about sensitivity to parameter values.

